# PathwayMatcher: proteoform-centric network construction enables fine-granularity multi-omics pathway mapping

**DOI:** 10.1101/375097

**Authors:** Luis Francisco Hernández Sánchez, Bram Burger, Carlos Horro, Antonio Fabregat, Stefan Johansson, Pål Rasmus Njølstad, Harald Barsnes, Henning Hermjakob, Marc Vaudel

## Abstract

**Background:** Mapping biomedical data to functional knowledge is an essential task in bioinformatics and can be achieved by querying identifiers, e.g. gene sets, in pathway knowledgebases. However, the isoform and post-translational modification states of proteins are lost when converting input and pathways into gene-centric lists.

**Findings:** Based on the Reactome knowledgebase, we built a network of protein-protein interactions accounting for the documented isoform and modification statuses of proteins. We then implemented a command line application called PathwayMatcher (github.com/PathwayAnalysisPlatform/PathwayMatcher) to query this network. PathwayMatcher supports multiple types of omics data as input, and outputs the possibly affected biochemical reactions, subnetworks, and pathways.

**Conclusions:** PathwayMatcher enables refining the network-representation of pathways by including isoform and post-translational modifications. The specificity of pathway analyses is hence adapted to different levels of granularity and it becomes possible to distinguish interactions between different forms of the same protein.

## Findings

In biomedicine, molecular pathways are used to infer the mechanisms underlying disease conditions and identify potential drug targets. Pathways are composed of series of biochemical reactions, of which the main participants are proteins, that together form a complex biological network [1]. Proteins can be found in various forms, referred to as proteoforms [2]. The different proteoforms that can be obtained from the same gene/protein depend on the individual genetic profiles, on sequence cleavage and folding, and on post-translational modification (PTM) states [3]. Proteoforms can carry PTMs at specific sites, conferring each proteoform unique structure and properties [4]. Notably, many pathway reactions can only occur if all or some of the proteins involved are in specific post-translational states.

However, when analyzing omics data, both input and pathways are summarized in a gene- or protein-centric manner, meaning that the different proteoforms and their reactions are grouped by gene name or protein accession number, and the fine-grained structure of the pathways is lost. One can therefore anticipate that proteoform-centric networks provide a rich new paradigm to study biological systems. But while gene networks have proven their ability to identify genes associated with diseases [5], networks of finer granularity remain largely unexplored.

Here, we present PathwayMatcher, an open-source standalone application that considers the isoform and PTM status when building protein networks and mapping omics data to pathways from the Reactome database. Reactome [6], is an open-source curated knowledgebase consolidating documented biochemical reactions categorized in hierarchical pathways, and notably includes isoform and PTM information for the proteins participating in reactions and pathways.

As an example of the complexity of hierarchical pathway information, we provide a graph representation of *Signaling by NOTCH2* from Reactome (**Figure 1**). This pathway is a sub-pathway of the pathways *Signaling by NOTCH* and *Signal Transduction*. It is composed of two sub-pathways (*NOTCH2 intracellular domain regulates transcription* and *NOTCH2 Activation and Transmission of Signal to the Nucleus*), comprising 32 and 54 reactions, yielding 28 and 141 edges, respectively. The 31 participants of the *Signaling by NOTCH2* pathway are also involved in reactions in other pathways, between themselves and with 2,055 other proteins, resulting in 6,525 external edges. Note that in this pathway, Cyclic AMP-responsive element-binding protein 1 (coded by *CREB1*) is phosphorylated at position 46 (labeled as *CERB1_P* in **Figure 1**) and Neurogenic locus notch homolog protein 2 (coded by *NOTCH2*), is found in three forms (unmodified and with two combinations of glycosylation, labeled as NOTCH2, NOTCH2_Gly1, and NOTCH2_Gly2, respectively).

**Figure 1:**
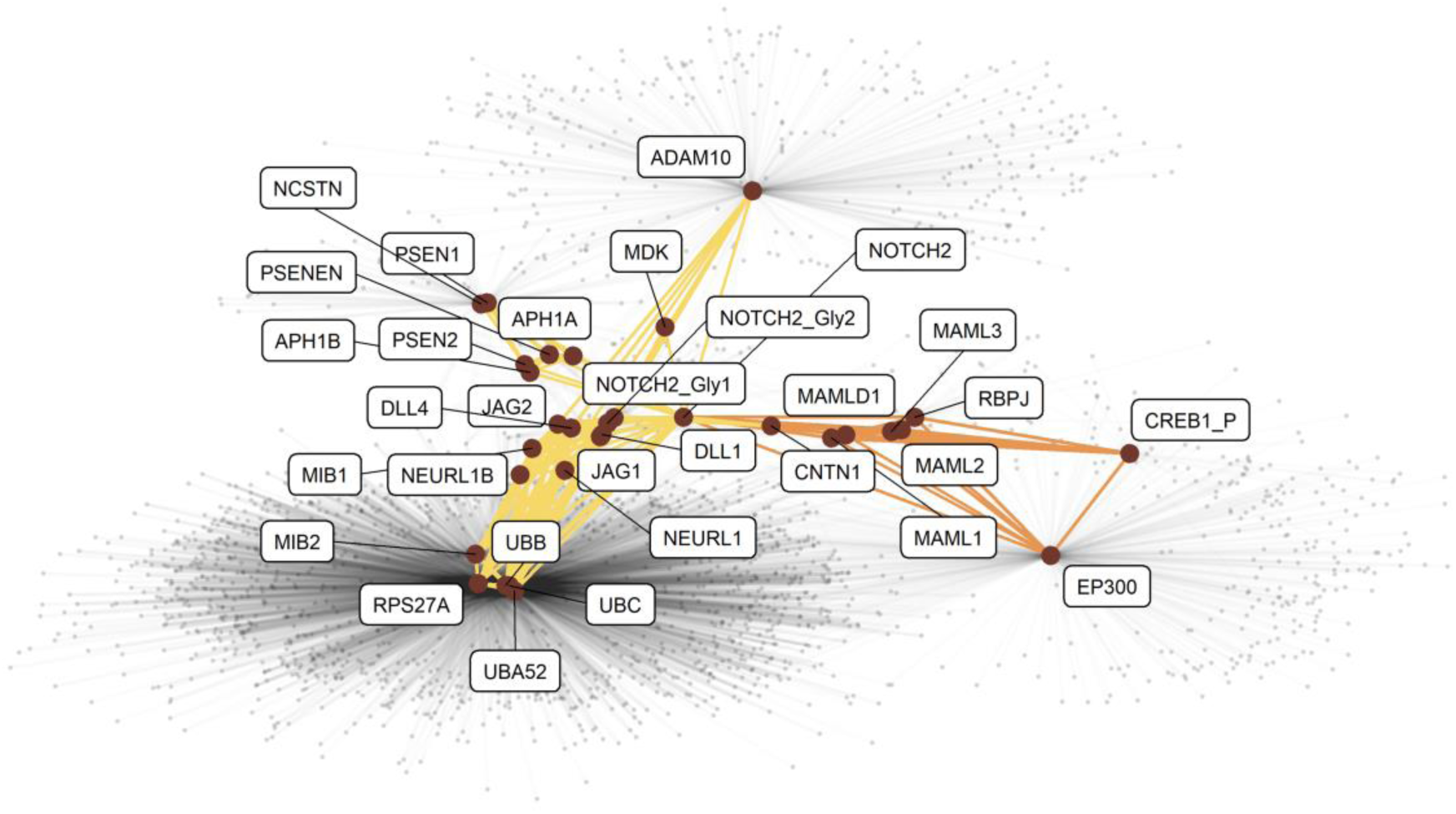
Graph representation of the Signaling by *NOTCH2* pathway as extracted from the Reactome database. Participating proteins are displayed as large dark red dots labeled with their canonical gene name. Post-translational modifications (PTMs) are indicated with suffixes in the label. A connection between two dots indicates a documented interaction between the two proteins in the given pathway. Connections belonging to the sub-pathways *NOTCH2* intracellular domain regulates transcription and *NOTCH2* Activation and Transmission of Signal to the Nucleus are displayed in orange and yellow, respectively. The interactions involving these proteins in other pathways are displayed with light gray connections in the background.

The amount of information available on reactions involving modified proteins has dramatically increased during the past two decades (**Figure 2**), with 3,947 and 5,631 publications indexed in Reactome (version 64 at time of writing) describing at least one reaction between modified proteins or between a modified and an unmodified protein, respectively. To harness this vast amount of knowledge, we built a network representation of pathways that we refer to as *proteoform-centric*, where protein isoforms with different sets of PTMs are represented with different nodes, in contrast to *gene-centric* networks, where one node is used per gene name or protein accession. In this representation, two proteoforms are connected if they participate in the same reaction. Note that proteoforms can participate in reactions both individually and as part of a set or complex. Furthermore, they can have four different roles: input, output, catalyst, or regulator.

**Figure 2:**
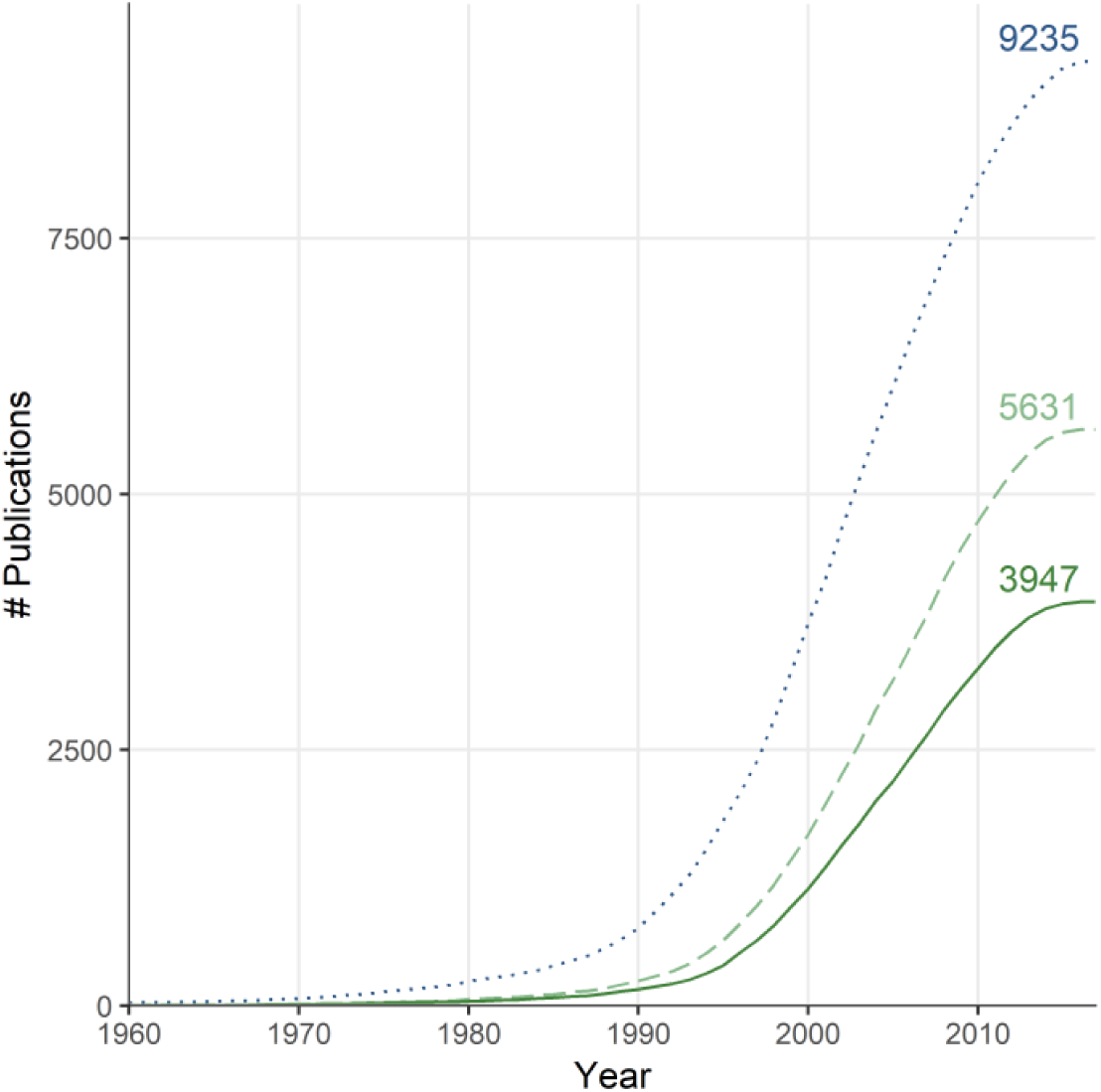
The number of publications indexed in Reactome documenting at least one reaction between two proteins with PTMs (solid dark green line), between one protein with PTMs and one without (dashed light green line), and two proteins without PTMs (dotted blue line), counting all publications with a year earlier than or equal to the x-axis value. The number of publications in each category at time of writing is indicated to the right.

The fundamental difference between gene- and proteoform-centric networks is illustrated in **Figure 3**, showing the graph representation of interactions with the protein *Cellular tumor antigen p53* (P04637) from the *TP53* gene. In a gene-centric paradigm (**Figure 3A**), 221 nodes are connected to a single node, making 220 connections; while in a proteoform-centric network (**Figure 3B**), 227 proteoforms connect to 23 proteoforms coded by *TP53* making 414 connections. Note that the proteoforms coded by *TP53* are themselves involved in reactions, making 24 *TP53-TP53* connections. In this example, the proteoform-centric network thus presents more nodes and connections than the gene-centric network, with visible structural differences in the network organization. We hypothesize that the proteoform-centric network paradigm depicted in Figure 3B provides a rich map that will enable navigating biomedical knowledge to a higher level of detail, to better assess the effect of perturbations, and identify drug targets more specifically.

**Figure 3:**
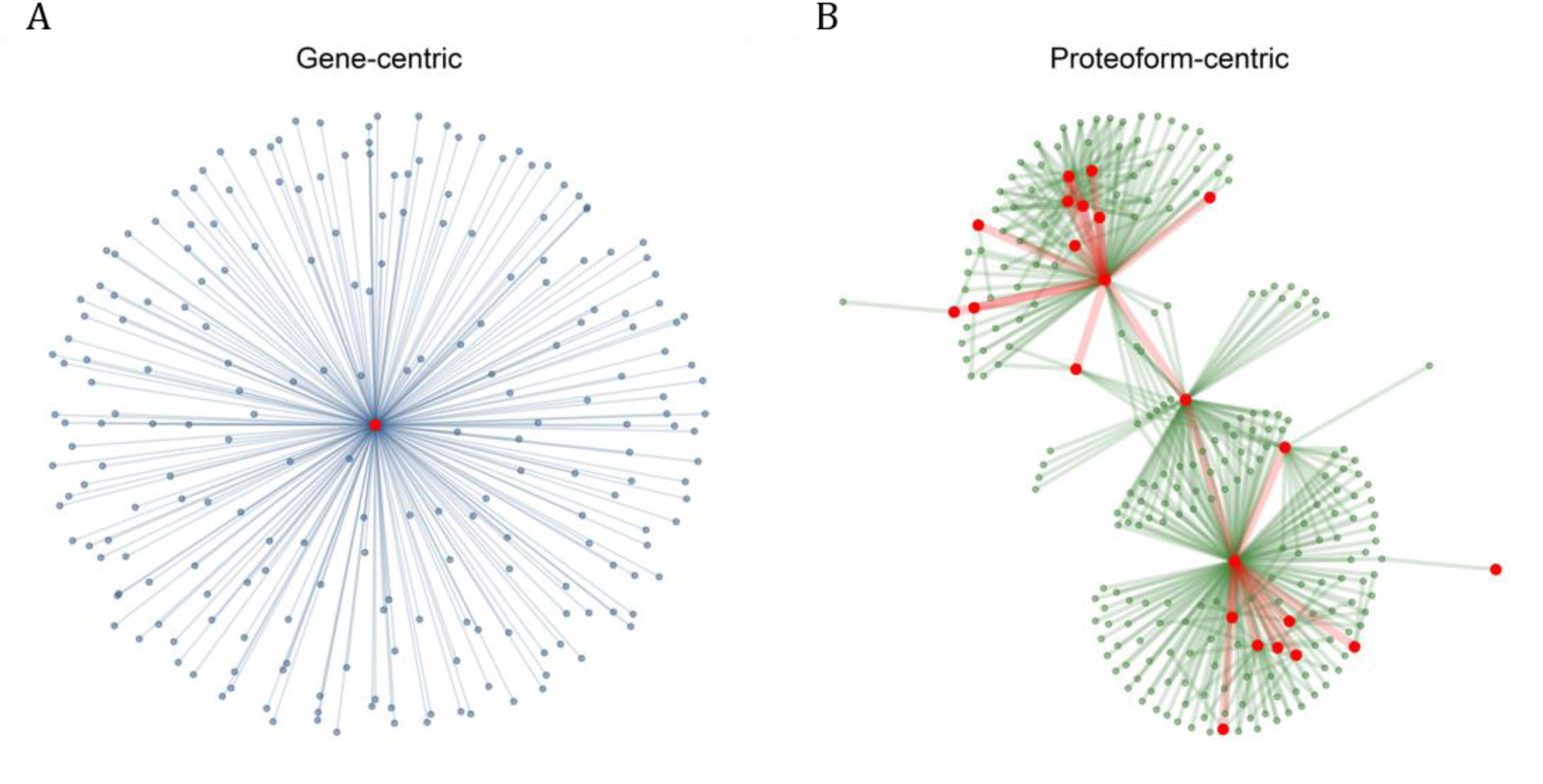
Gene-centric versus protein-centric representation. (A) Graph representation of the genes involved in reactions (through their corresponding proteins) with (the corresponding proteins of) *TP53*, with a single node per gene. *TP53* is represented with a large red dot at the center and genes coding proteins involved in reactions with *TP53* are represented with smaller blue dots at the periphery connected to the *TP53* gene with blue lines. (B) Graph representation of the proteins involved in a reaction with gene products of *TP53*, distinguishing isoforms and post-translationally modified proteins as different proteoforms. The proteoforms coded by *TP53* and the proteoforms involved in a reaction with them are represented with large red and small green dots, respectively. The connections between proteoforms coded by *TP53* are displayed with thick red lines and connections with other proteoforms with thin green lines.

PathwayMatcher allows the user to tune the granularity of the network representation of pathways by representing nodes as (i) gene names, (ii) protein accession numbers, or (iii) proteoforms, and supports the mapping of multiple types of omics data: (i) genetic variants, genes, (iii) proteins, (iv) peptides, and (v) proteoforms. Genetic variants are mapped to proteins using the Ensembl Variant Effect Predictor [7], gene names are mapped to proteins using the UniProt identifier mapping [8], and peptides are mapped to proteins using PeptideMapper [9]. If a peptide maps to different proteins, all possible proteins are considered for the search and protein inference must be conducted *a posteriori* [10]. If peptides are modified, they are mapped to the proteoforms presenting compatible PTM sets. Proteins are mapped to the pathway network using their accession, while proteoforms are mapped by comparing their protein accession, isoform number, and PTM set. A schematic representation of the PathwayMatcher matching procedure is shown in **Figure 4**. More details on the mapping procedure, formats, and settings can be found in the methods section and in the online documentation (github.com/PathwayAnalysisPlatform/PathwayMatcher/wiki).

**Figure 4:**
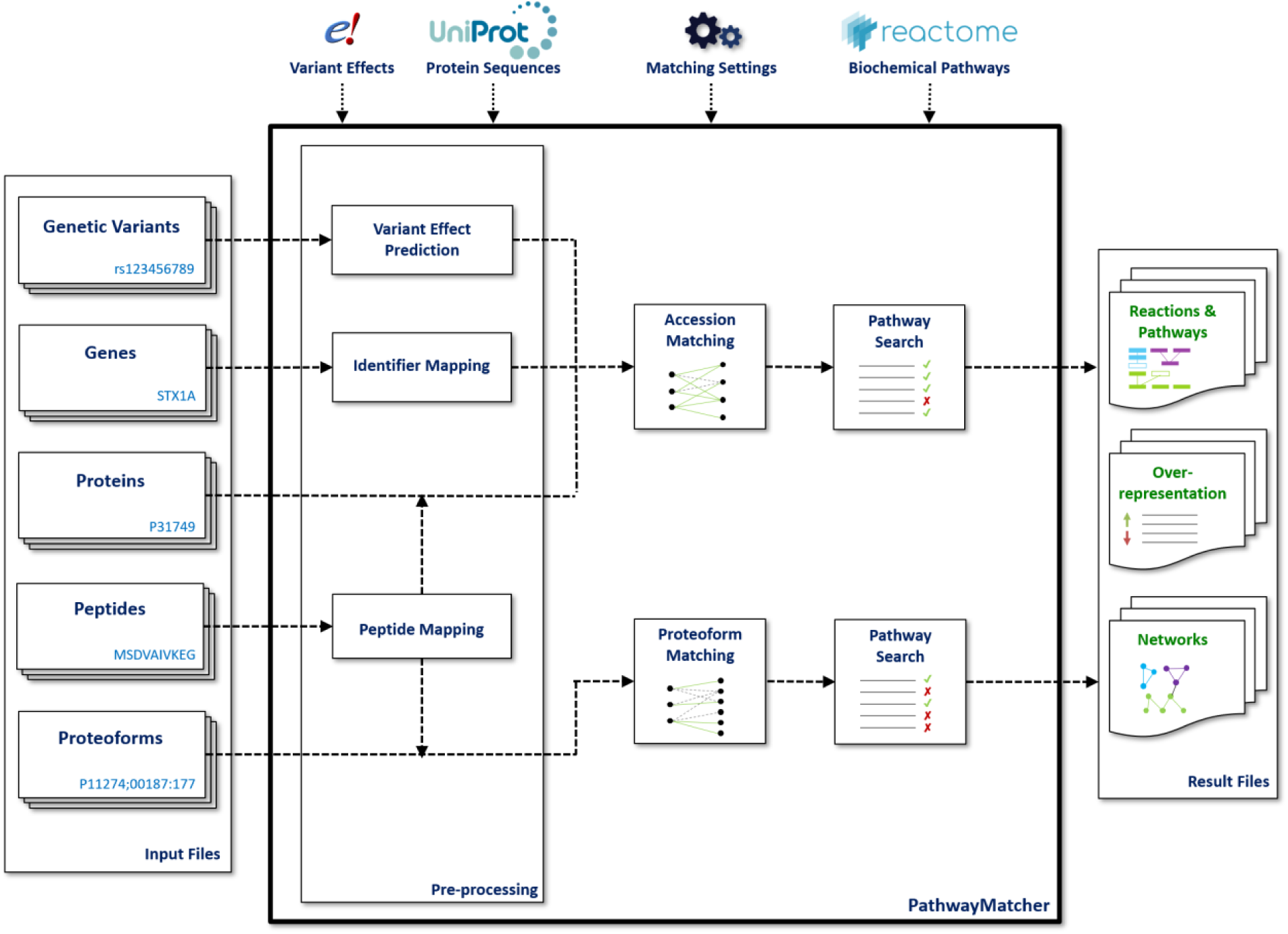
Schematic representation of the PathwayMatcher matching procedure. Input of various types is modelized as sets of proteins or proteoforms based on the annotation of isoforms and PTMs. Proteins and proteoforms are then mapped to Reactome based on user settings. Matched reactions and pathways, the results of an over-representation analysis, and sub-networks generated from the input are exported as text files.

PathwayMatcher produces three types of output: (i) the result of the matching, listing all possible reactions and pathways linked to the input; (ii) the results of an over-representation analysis; and (iii) networks in relationship with the input. The over-representation analysis is performed on the pathways matching and follows the first generation of pathway analysis methods [11], *i.e.* a *p*-value for each pathway in the reference database is calculated using a binomial distribution followed by Benjamini-Hochberg correction [12] (in a similar way as performed by the Reactome online analysis tool [13]). If the input can be mapped to proteoforms, the over-representation analysis is conducted using a proteoform-centric representation of pathways, using proteins otherwise. The exported networks represent the internal and external connections that can be drawn from the input, where internal connections connect two nodes from the input list, and external connections one node from the input list to any node not in the input. The user can select to export these networks using nodes defined as genes, proteins, or proteoforms. Connections between nodes in the network are annotated with information on whether they participate as complex or set, and their role in the reaction.

As displayed in **Figure 5A**, 68% of the pathways present at least one proteoform-specific participant, *i.e.* with isoform or PTM annotation. The number of pathways containing a given gene product or proteoform is displayed in **Figure 5B**, showing how using proteoforms allows distinguishing pathways more specifically than genes, with a median of four pathways matched per proteoform compared to eleven pathways per gene. When the input can be mapped to proteoforms, PathwayMatcher can restrict the search for reactions and pathways to those that specifically involve proteins in the desired form, hence reducing the number of possible connections for a given node in the resulting network. Conversely, the proteoform-centric network representation allows identifying interactions between multiple proteoforms originating from the same gene or protein, resulting in new connections compared to a gene-centric representation.

**Figure 5:**
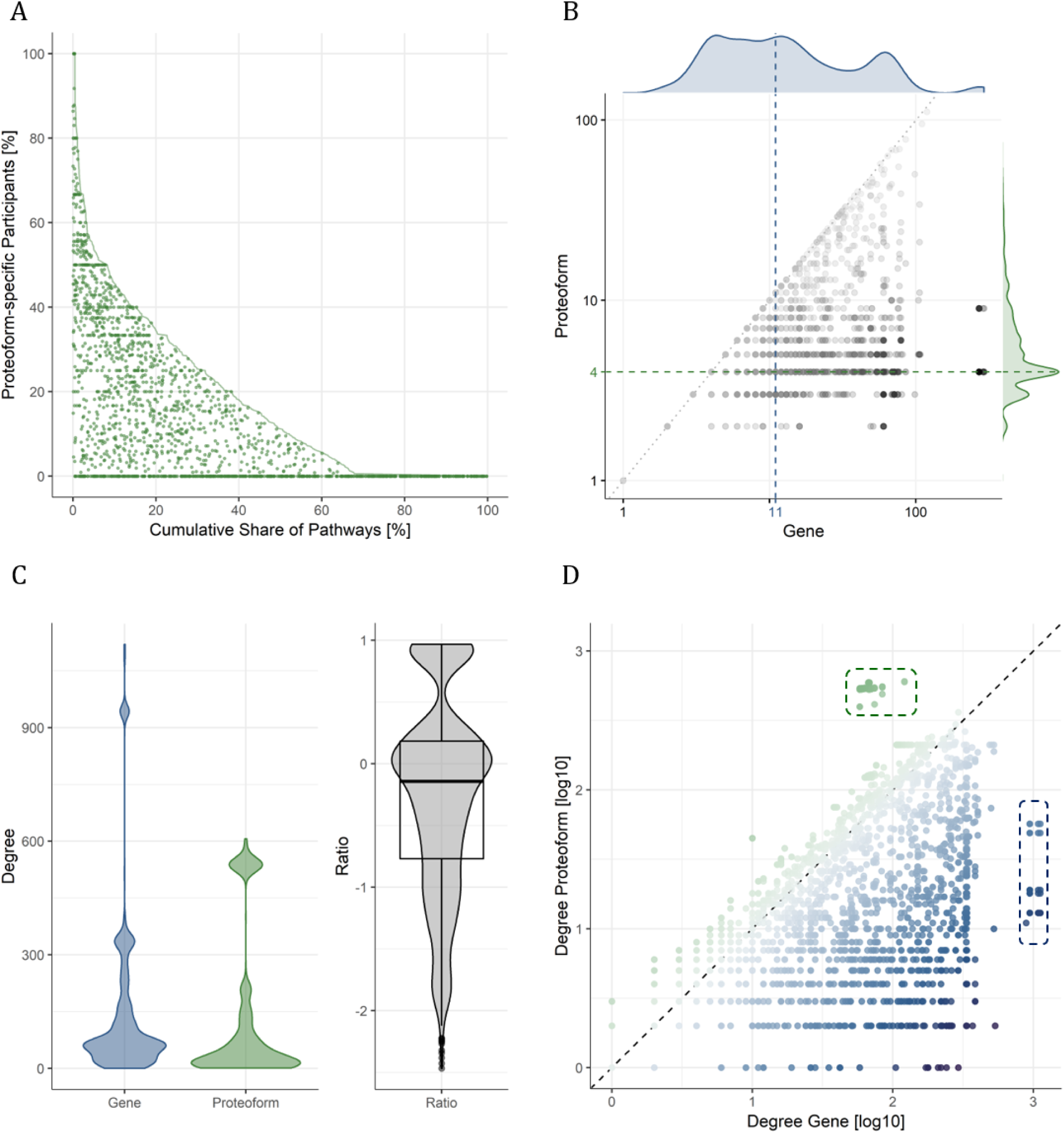
Prevalence of proteoforms in pathways. (A) The share of proteoform-specific participants in a pathway, i.e. proteins that are annotated with isoform and/or PTM information, is plotted against the cumulative share of pathways, going from the highest share of proteoforms to the lowest. The cumulative share of pathways is displayed with a solid green line. The share of proteoform-specific participants in each pathway is plotted with a green dot with a jitter on the x-axis between zero and the solid line. (B) For all proteoform-specific participants, the number of pathways mapped using the proteoform versus gene is plotted in black. The density of the number of pathways mapped are indicated at the top (blue) and right (green) for gene and proteoform matching, respectively. The median number of pathways mapped is indicated with dashed lines. (C) Violin plots of the degree, i.e. number of connections, for the proteoform-specific participants in a gene-centric (blue) or proteoform-centric network (green) are plotted to the left; violin and box plots of the ratio of degrees, proteoform over gene, are plotted to the right (black). (D) The degree of the proteoform-specific participants in the proteoform-centric network is plotted against the degree in the gene-centric network. Note that base 10 logarithmic scales are used on both axes. Dots are colored with a blue-grey-green gradient corresponding to the ratio in C. Outliers of high degree in the gene-centric but not in the proteoform-centric network are indicated with blue dashes to the right. Outliers of high degree in the proteoform-centric but not in the gene-centric network are indicated with green dashes to the top.

**Figure 5C** shows that the number of connections per proteoform is lower than the number of connections for the respective gene for most proteoforms, varying from 300-fold decrease to 10-fold increase. Interestingly, plotting the number of connections of a proteoform in gene-centric or proteoform-centric networks shows that the largest gene-centric hubs, corresponding to five genes, decompose into 127 proteoforms that do not outlie the distribution of the number of connections in the proteoform network (**Figure 5D**). Conversely, a group of 484 densely connected outliers emerges from 44 genes.

Through its paradigm-shift, PathwayMatcher hence provides a fine-grained representation of pathways for the analysis of omics data. However, this comes at the cost of increased complexity: gene-centric networks comprise a limited number of nodes, approximately 20,000 for humans, whereas in a proteoform-centric paradigm, the human network is expected to have several million nodes [14]. With the current version of Reactome, building the gene- and proteoform-centric networks results in 9,759 and 12,775 nodes with 443,229 and 672,047 connections, respectively. We classified the nodes into two categories, canonical or specific gene products, depending on whether or not they represent the unmodified canonical isoform of a protein according to UniProt. Within the proteoform network, 432,169 connections between 9,694 nodes link two canonical gene products, 95,539 connections between 7,734 nodes involved one canonical and one specific gene product, and 2,806 nodes with 144,339 connections involved two specific gene products.

In addition to the increased size of the underlying network, matching proteoforms requires comparing isoforms and sets of modifications, possibly with tolerance and wildcards for the modification definition and localization, which is computationally much more intensive than simply comparing identifiers. **Figure 6** shows the performance of PathwayMatcher benchmarked against public data sets of (A) genetic variants, (B) proteins, (C) peptides, and (D) proteoforms.

**Figure 6:**
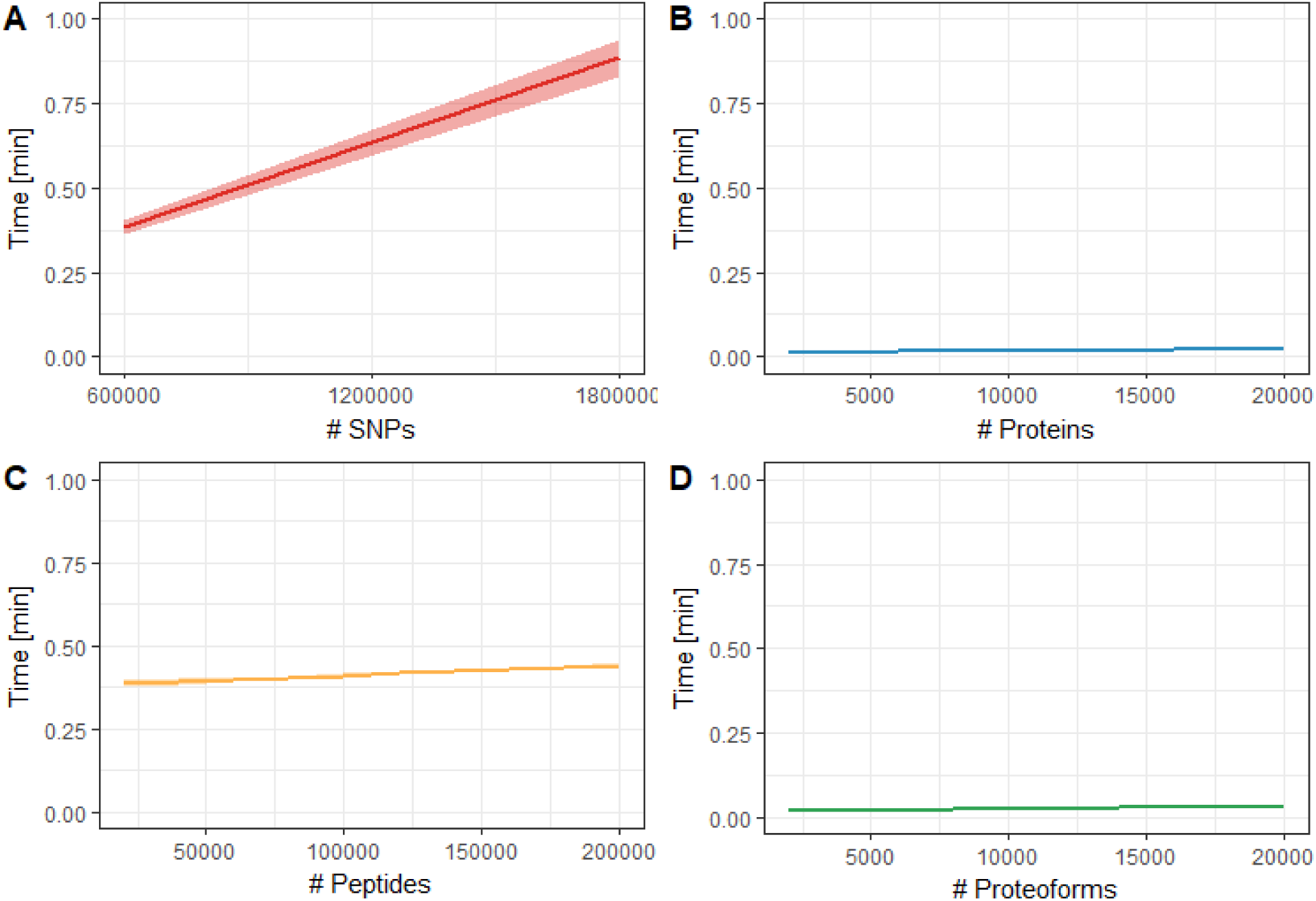
Performance of PathwayMatcher using (A) genetic variants as single-nucleotide polymorphisms (SNPs), (B) proteins, (C) peptides and (D) proteoforms. Performance in minutes is plotted against input size. The mean is displayed as a solid line and the 95% range as a ribbon (only visible in (A) due to the high reproducibility in other cases).

For the proteins and proteoforms, the processing time increased linearly related to the query size with a small slope, making it possible to search all available proteins within a few seconds. As expected, protein identifiers provided the fastest response time, while proteoforms were the second fastest. Mapping peptides took approximately 30 seconds more, corresponding to the indexing time of the protein sequences database by PeptideMapper [9], after which the time increased linearly in a similar fashion as for proteins. For the genetic variants, an extra mapping step is required to map possibly affected proteins, adding additional computing time. The overall mapping time for a million single-nucleotide polymorphisms (SNPs) was less than a minute, which is acceptable compared to the other steps of a variant analysis pipeline. Note that the processing time was very reproducible across runs, where minor variation is only noticeable using genetic variants, resulting in very thin ribbons in **Figure 6B-D**.

In conclusion, PathwayMatcher is a versatile application enabling the mapping of several types of omics data to pathways in reasonable time and can readily be included in bioinformatic workflows. Thanks to the fine-grained information available in Reactome, PathwayMatcher supports refining the pathway representation to the level of proteoforms. To date, only a fraction of the several million expected proteoforms [14] have annotated interactions, but as the understanding of protein interactions continues to increase, and the ability to identify and characterize them in samples progresses, proteoform-centric networks will surely become of prime importance in biomedical studies. Notably, the effect of genetic variation on genes, transcripts, and proteins is currently only partially resolved for a fraction of the genome. The rapid development of this field will make it possible to identify biological functions affected by variants within the human network. Refining its representation to the level of proteoforms will allow pinpointing more precisely reactions and pathways, and hence increase our ability to understand biological mechanisms and potentially identify druggable targets.

## Methods

### Implementation

PathwayMatcher is implemented in Java 8.0.

### Availability

PathwayMatcher is freely available at github.com/PathwayAnalysisPlatform/PathwayMatcher under the permissive Apache 2.0 license. It is also possible to use PathwayMatcher as a Docker image: hub.docker.com/r/lfhs/pathwaymatcher. PathwayMatcher can be obtained from the Bioconda channel of the Conda [15] package manager at bioconda.github.io/recipes/pathwaymatcher/README.html. Finally, PathwayMatcher is available as a Galaxy [16] tool in the Galaxy ToolShed [17] at toolshed.g2.bx.psu.edu/view/galaxyp/reactome_pathwaymatcher where it can be readily integrated into analysis workflows. PathwayMatcher has also been installed into the public European Galaxy instance, usegalaxy.eu, making it possible to use the application without requiring any local configuration and just providing valid input files and options. The complete URL for the online tool is: https://usegalaxy.eu/?tool_id=toolshed.g2.bx.psu.edu%2Frepos%2Fgalaxyp%2Freactome_pathwaymatcher%2Freactome_pathwaymatcher

Upon installation, PathwayMatcher can be used from the command line to query Reactome using various types of omics data. Either the “.jar” file is run directly using Java or the Docker image is instantiated to a container. Detailed information on implementation, installation, usage and format specifications is available in the online documentation at github.com/PathwayAnalysisPlatform/PathwayMatcher/wiki.

### Input and Output

Detailed and updated documentation of the input and output can be found in the online documentation at github.com/PathwayAnalysisPlatform/PathwayMatcher/wiki.

As schematized in **Figure 7**, a simple representation is used for proteoforms: (i) the UniProt protein accession and (ii) the set of PTMs separated by a semicolon ‘;’. The protein accession can include the isoform number specified with a dash ‘-‘. The PTM set contains each PTM separated by a comma ‘,’. Each PTM is specified using a modification identifier and a site, separated by a colon ‘:’.

**Figure 7:**
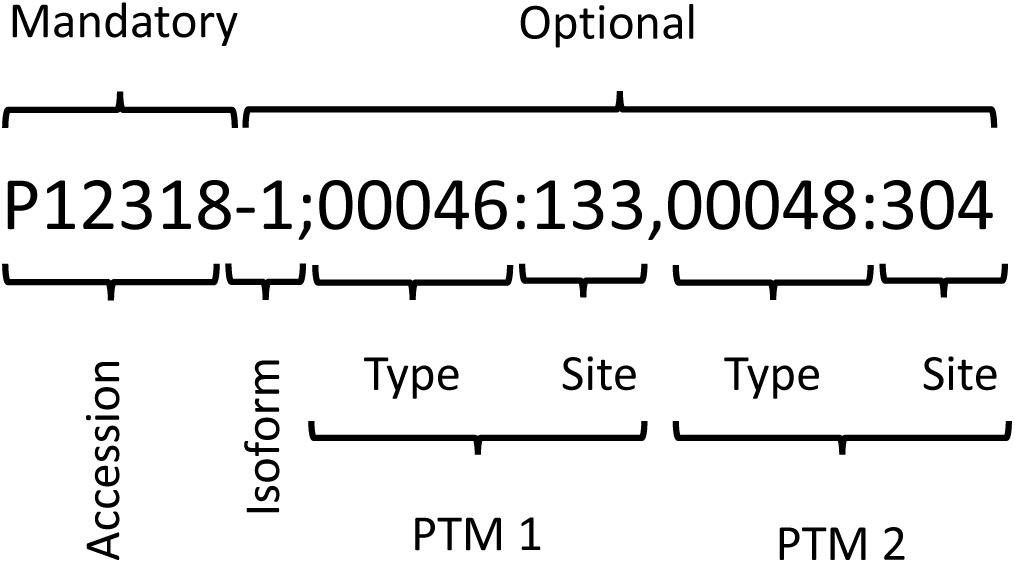
Example of identifier for a proteoform, composed of a protein accession, an isoform number, and a set of PTMs.

Note that the order of PTMs do not affect the search. The PTM identifier is a five digit identifier from the PSI-MOD Protein Modification [18]. The site is an integer specifying the 1-based index of the modified amino acid on the sequence as defined by UniProt.

It is common to write the identifiers for the PTM types with the prefix ‘MOD:’ before the five digits of the ontology term. PathwayMatcher also allows the user to write the identifier without the prefix. PathwayMatcher also allows querying all proteoforms modified at a given site using the ‘00000’ wild card for modification type. Note that the modification site is mandatory, but a tolerance window can be set, as detailed in the “Proteoform matching” section.

### Post-translational modifications in the Reactome data model

The Reactome object model specifies physical entities, *e.g.* complexes, proteins and small molecules, and proteins are annotated using unique identifiers. These entities participate in reactions in specific cellular compartments. They can also be connected to multiple instances of *Translational Modification* objects containing a specific coordinate on the protein sequence and an identifier following the PSI-MOD ontology [18]. The portion of physical entities referring to proteins are associated to other class of objects as reference entities, which contain protein annotations in external databases such as UniProt [19]. Therefore, a proteoform is represented as a physical entity associated to a set of modifications for specific processes at specific subcellular location. 127 different protein modifications are annotated in Reactome for humans, of which **Figure 8** displays the most frequent.

**Figure 8:**
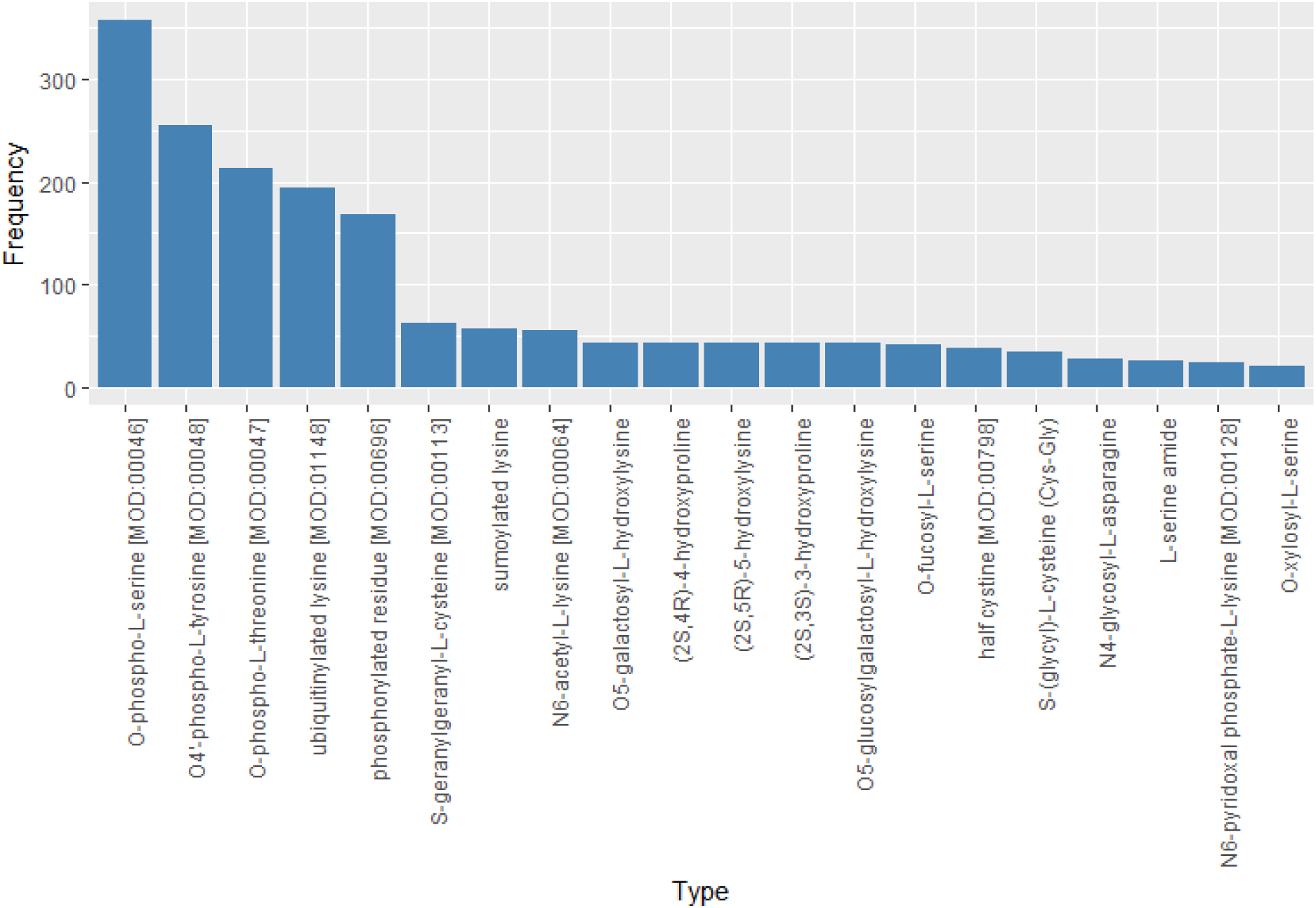
Prevalence of the different PTM annotations in Reactome. PTM labels are extracted from the Reactome database and the number of proteins annotated with the PTM is displayed for each label. If a protein is carrying multiple instances of the PTM, the PTM is counted only once.

### Proteoform matching

Searching pathways using gene names or protein accessions solely requires mapping a string of characters between the input and the knowledgebase. In order to map the proteoforms to reactions and pathways, it is necessary to decide if the proteoforms in the input are equivalent to the proteoforms annotated in the Reactome database, taking into account the protein accession, isoform information, and the set of PTMs. Two proteoforms can have all, some or none of these elements in common.

We defined a set of criteria to compare two proteoforms, one from the input and one from the reference database and decide whether they are equivalent to each other. The matching types defined for PathwayMatcher are: a) S*uperset*, b) S*ubset*, c) *One*, and d) *Strict*.

#### a) Superset (with and without PTM types)

The set of input PTMs is a superset of the reference PTMs set. This includes the command line arguments: -m **superset** or -m **superset_no_types**.

- The UniProt accession is the same
- The isoform is the same; either:
  - Both have an isoform specified. Ex: P31749-3
  - Or both refer to the default one. Ex: P31749
- The PTMs:
  - The input contains ALL the reference PTMs or more (input is superset or equal). Each reference PTM must have a matching input PTM. Some input PTMs may not have a matching reference PTM.
- A PTM matches if these two requirements are true:
  - The types match:
    - If chosen **superset** then types should be equal.
    - If chosen **superset_no_types** the type is not considered.
  - The coordinates match if any of the following is true:
    - Both are known identical (positive integer) coordinates.
    - Both are known different (positive integer) coordinates, but the absolute difference between the two coordinates is less than or equal to a user-defined margin (‘range’ option in command line).
    - One of the coordinates is unknown (null, empty, ?, −1).

#### b) Subset (with and without PTM types)

The set of input PTMs is a subset of the reference PTMs set. This includes the command line arguments: -m **subset** or -m **subset_no_types**.

- The UniProt accession is the same.
- The isoform is the same; either:
  - Both have an isoform specified. Ex: P31749-3
  - Or both refer to the default one. Ex: P31749
- The PTMs:
  - Each input PTM must have a matching reference PTM. Some reference PTMs may not have a matching input PTM.
- A PTM matches if these two requirements are true:
  - Types match, i.e.:
    - If chosen **subset** (then types must be equal), or
    - If chosen **subset_no_types** (type is not considered)
  - The coordinates match if any of the following is true:
    - Both are known identical (positive integer) coordinates.
    - Both are known different (positive integer) coordinates but the absolute difference between the two coordinates is less than or equal to a user-defined margin (‘range’ option in command line).
    - One of the coordinates is unknown (null, empty, ?, −1).

#### c) One (with and without PTM types)

At least one input PTM matches. This includes the command line arguments: -m **one** or –m **one_no_types**.

- The UniProt accession is the same.
- The isoform is the same; either:
  - Both have an isoform specified. Ex: P31749-3
  - Or both refer to the default one. Ex: P31749
- The PTMs:
  - At least one input PTM must have a matching reference PTM.
- A PTM matches if these two requirements are true:
  - The types match:
    - If chosen one (then types should be equal), or
    - If chosen one_no_types (type is not considered)
  - The coordinates match if any of the following it true:
    - Both are known identical (positive integer) coordinates.
    - Both are known different (positive integer) coordinates, but the absolute difference between the two coordinates is less than or equal to a user-defined margin (‘range’ option in command line).
    - One of the coordinates is unknown (null, empty, ?, −1).

#### d) Strict

Proteoforms must match exactly in all the attributes.

- The UniProt accession is the same.
- The isoforms are the same; either:
  - Both have an isoform specified. Ex: P31749-3
  - Or both refer to the default one. Ex: P31749
- The PTMs have the same elements:
  - The reference PTM set and the input PTM set have the same size.
  - Each reference PTM has a matching input PTM.
- A PTM matches if:
  - Types are the same.
  - Coordinates are the same:
    - In case they are numbers, they should be equal
    - In case they are null, then both should be null.

Extra considerations:

- Negative, zero or floating-point values are invalid as sequence coordinates in the input.
- We accept only PSI-MOD ontology modification types.
- The margin to compare the coordinates should be set as an unsigned integer.

**Table 1** shows examples of PTM coordinates matching. The letter *k* represents any positive integer. It compares a PTM coordinate in an input PTM with a PTM coordinate in a reference PTM.

**Table 1:** Post-translational modification coordinates comparison criteria.

| Input | Reference | Margin | Matched | Comment |
| --- | --- | --- | --- | --- |
| 17 | 17 | 0 | Yes | Equal |
| 16 | 17 | 0 | No | Out of margin |
| 18 | 17 | 0 | No | Out of margin |
| 7 | 13 | 5 | No | Out of margin |
| 8 | 13 | 5 | Yes | In margin |
| 9 | 13 | 5 | Yes | In margin |
| 17 | 13 | 5 | Yes | In margin |
| 18 | 13 | 5 | Yes | In margin |
| 19 | 13 | 5 | No | Out of margin |
| 0 | 2 | 5 | No | Input in margin but not valid |
| -1 | 2 | 5 | No | Input in margin but negative |
| ?, empty, null | Positive integer | k | Yes | Input is less specific |
| Positive integer | ?, empty, null, -1 | k | Yes | Input is more specific |
| ?, empty, null | ?, empty, null, -1 | k | Yes | Equally unspecific |
| Negative int, zero | Any | k | No | Negative or zero input are invalid |

### Mapping omics data to pathways

The input is mapped to proteins or proteoforms to find the reactions where the input entities are participants (**Figure 9**). The input is mapped to proteins when data types without PTMs or specific translation products are specified; otherwise a mapping to proteoforms is used. When one type of data yields multiple results due to ambiguity, e.g. a SNP or peptide mapping multiple proteins, all the possibilities are included in the search entities.

**Figure 9:**
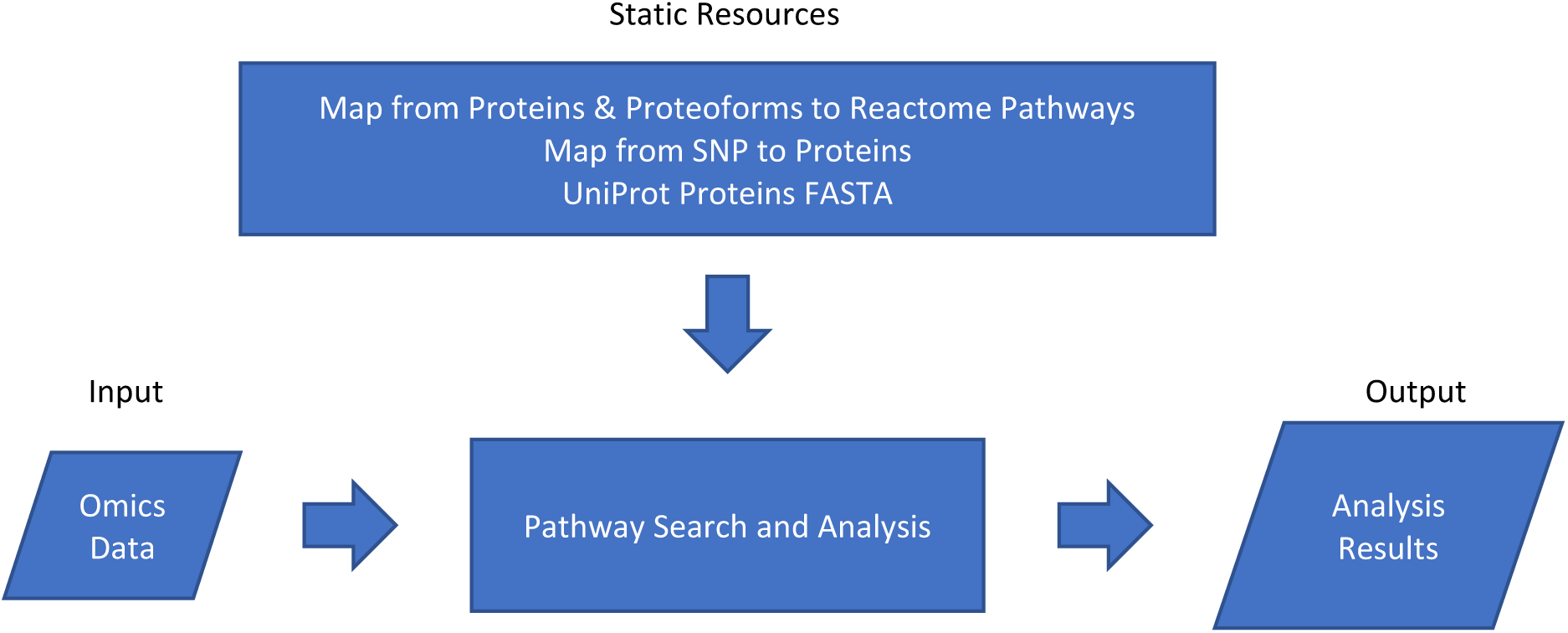
PathwayMatcher general overview. The program takes the user input in the form of omics data files and the reference pathways from the database as input. It then executes the search and analysis algorithm to create a resulting list of output files.

When a list of SNPs is provided, mapping from the Ensembl Variant Effect Predictor (VEP) [7] is used to find the possibly affected proteins. When peptides are provided, their sequence is mapped to UniProt protein identifiers [8] using PeptideMapper [9] and possible proteoforms are constructed. When proteins or proteoforms are available, PathwayMatcher maps them to reactions and pathways using data structures embedded in the PathwayMatcher jar file. These data structures are extracted from the Reactome Neo4j graph database (neo4j.com) and serialized. The code for extraction of the relationships from proteins to pathways is available at github.com/PathwayAnalysisPlatform/Extractor.

Proteins in Reactome are defined according to UniProt following a gene-centric paradigm. From all reactions in Reactome, 9,734 involve two proteins, participating in 2,208 human pathways [1] (version 64 at time of writing). Using additional information from Reactome on the post-translational state required for a protein to participate in a reaction, PathwayMatcher allows matching proteoforms to reactions and pathways. **Tables 2 and 3** respectively list the proteins and proteoforms that are participating in the highest number of pathways.

**Table 2:** Human proteins participating in the highest number of pathways in Reactome. Note that the Reactions Mapped column shows the number of reactions that are part of the mapped pathways. A protein may participate in a reaction that was not assigned to a pathway, and a reaction can be included in multiple pathways.

|  | Gene | Protein name | Reactions<br>Mapped | Pathways<br>Mapped |
| --- | --- | --- | --- | --- |
| P62979 | RPS27A | Ubiquitin-40S ribosomal protein S27a | 306 | 292 |
| P62987 | UBA52 | Ubiquitin-60S ribosomal protein L40 | 299 | 288 |
| P0CG47 | UBB | Polyubiquitin-B | 279 | 270 |
| P0CG48 | UBC | Polyubiquitin-C | 279 | 270 |
| P62993 | GRB2 | Growth factor receptor-bound protein 2 | 259 | 144 |
| P28482 | MAPK1 | Mitogen-activated protein kinase 1 | 70 | 119 |
| P30153 | PPP2R1A | Serine/threonine-protein phosphatase 2A 65 kDa regulatory subunit A alpha isoform | 83 | 116 |
| P01112 | HRAS | GTPase HRas | 89 | 112 |
| P01116 | KRAS | GTPase KRas | 87 | 108 |
| Q07889 | SOS1 | Son of sevenless homolog 1 | 118 | 107 |

**Table 3:** Human proteoforms participating in the highest number of pathways in Reactome.

|  | Gene | Protein name | PTMs | Reactions<br>Mapped | Pathways<br>Mapped |
| --- | --- | --- | --- | --- | --- |
| P28482 | MAPK1 | Mitogen-activated protein kinase 1 | [00047:185, 000<br>48:187] | 48 | 111 |
| P27361 | MAPK3 | Mitogen-activated protein kinase 3 | [00047:202, 000<br>48:204] | 43 | 95 |
| P31749 | AKT1 | RAC-alpha serine/threonine-protein<br>kinase ecNumber2.7.11.1/ecNumber | [00046:473, 000<br>47:308] | 51 | 78 |
| P31751 | AKT2 | RAC-beta serine/threonine-protein<br>kinase ecNumber2.7.11.1/ecNumber | [00046:474, 000<br>47:309] | 33 | 69 |
| Q9GZV9 | FGF23 | Fibroblast growth factor 23 | [00164:178] | 152 | 68 |
| P16220 | CREB1 | Cyclic AMP-responsive element-binding protein 1 | [00046:133] | 13 | 68 |
| Q16539 | MAPK14 | Mitogen-activated protein kinase 14 | [00047:180, 000<br>48:182] | 16 | 63 |
| Q15759 | MAPK11 | Mitogen-activated protein kinase 11 | [00047:180, 000<br>48:182] | 15 | 57 |
| P19174 | PLCG1 | 1-phosphatidylinositol 4,5-bisphosphate phosphodiesterase<br>gamma-1<br>ecNumber3.1.4.11/ecNumber | [00048:472, 000<br>48:771, 00048:7<br>83, 00048:1253] | 38 | 55 |
| Q8WU20 | FRS2 | Fibroblast growth factor receptor<br>substrate 2 | [00048:196, 000<br>48:306, 00048:3<br>49, 00048:392, 0<br>0048:436, 00048<br>:471] | 94 | 53 |

Note that PathwayMatcher maps experimental data to pathways in a systematic and unbiased fashion. This means that it collects all pathways containing at least one of the participant proteins or proteoforms of the input data and does not perform any filtering or biological inference. Therefore, it attempts at minimizing the prevalence of false negatives by considering all the possible pathways annotated in the reference database. It can however not control for missing annotation, *i.e.* what is not annotated in the knowledgebase is not considered to be happening.

### Over-representation analysis

The matching of each entity to a given pathway is modelled as a Bernoulli trial with two possible outcomes: success or failure, depending on whether the protein or proteoform is a participant of a reaction in the pathway. Trials are considered independent from each other, meaning that the outcome of previous trials does not affect the next. Finally, the probability of success is calculated by the proportion of choosing a protein in a pathway over the total number of possible proteins, therefore the probability is constant over all trials.

First, we search all the input entities (proteins or proteoforms) across all the pathways and count how many of them were found in each pathway. The number of entities found in a pathway is taken as the number of successful trials. Then, with the binomial probability distribution, we calculate how likely it would be to get a result equal to or more extreme than the current result (the same number or more proteins or proteoforms in the pathway), given that the input (proteins or proteoforms) were randomly selected [11].

This is done using the cumulative distribution function for the binomial distribution, which calculates the probability of getting at most *k* successes out of *n* trials, with a probability *p* ∈ [0,1], where *X* is a random variable following the binomial distribution, as detailed in Equation 1.

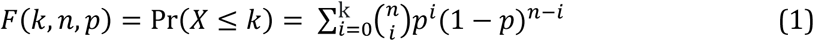

For each pathway, *p* is set to the ratio between the number of total proteins or proteoforms in the pathway and the total possible entities in the database, *n* is the number of proteins or proteoforms in the input sample, *k* is the number of proteins successfully mapped in the pathway, *X* is the number of entities found in the current pathway after the search.

Finally, given that the *p*-value requires the calculation of the probability of an equal or more extreme result, we use the complement of Equation 1 to calculate the probability of getting at least *k* successful trials out of *n* as stated in Equation 2.

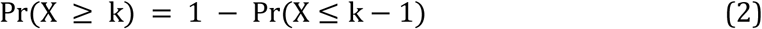

The calculations for proteins or proteoforms are similar, but are performed separately depending on the input. If the input consists of protein accessions, the number of participants is calculated by only considering proteins. On the other hand, for the proteoform input, the number of entities in the pathways and the database are the participant proteoforms.

Please note that the over-representation analysis is included as a simple analysis to identify the most covered pathways. We recommend however that users rather interpret the results of the mapping and the networks using the systems biology method that best suits the experiment and biomedical context. PathwayMatcher is developed to be a hypothesis generation tool, helping to navigating large datasets and guide experiments. It is not a validation or mechanism inference tool.

### Performance Benchmark

The performance of PathwayMatcher was evaluated using data sets of different sizes obtained from sampling publicly available resources:

- Proteins: human complement of the UniProtKB/Swiss-Prot database (release 2017_10).
- Peptides: ProteomeTools [20] as available in PRIDE [21], dataset PXD004732, release date 23/01/2017.
- Genetic variants: variants from the human assembly GRCh37.p13.
- Proteoforms: annotated proteoforms in Reactome Graph database version 62.

Performance testing was done using a standard desktop computer (Intel^®^ Core™ i7-6600U CPU @ 2.60GHz with 2 cores using 64-bit Windows 10 with Java SE 1.8.0_144 on SSD). Details and code are available at github.com/PathwayAnalysisPlatform/PathwayMatcher/wiki/Test-datasets.

### Metrics and Figures

The metrics presented in this manuscript were obtained by querying the Reactome graph database directly [22]. The queries used can be found in the online documentation at: github.com/PathwayAnalysisPlatform/PathwayMatcher/blob/master/docs/queriesForStatistics.md The figures in this manuscript were built in R version 3.4.1 (2017-06-30) - “Single Candle” (r-project.org) using the following packages: ggplot2, ggrepel, igraph, scico, grid, and gtable. The R scripts used to build the figures are available in the tool repository at: github.com/PathwayAnalysisPlatform/PathwayMatcher/tree/master/docs/figures/scripts

## Availability of supporting source code and requirements

**Project name:** PathwayMatcher

**Project home page:** github.com/PathwayAnalysisPlatform/PathwayMatcher

**Operating system(s):** Platform independent

**Programming language:** Java **Other requirements: License:** Apache 2.0

**RRID:** SCR_016759

## Declarations

**List of abbreviations**
PTM: Post-translational modification

### Ethics approval and consent to participate

Not applicable

### Consent for publication

Not applicable

### Authors’ contributions

LFHS did all of the programming, testing, documentation, and wrote the manuscript. BB and CH contributed with programming, testing, documentation, ideas, and manuscript writing. AF, SJ, and PRN contributed with ideas and manuscript writing. HB contributed with ideas, supervised the work, and wrote the manuscript. HH contributed with project design, manuscript writing, and supervised the work. MV contributed with project design, programming, documentation, testing, supervised the work, and wrote the manuscript. All authors participated in the preparation of the manuscript.

## Acknowledgements

The authors thank the Reactome curators for their massive curation effort and the guidance in interpreting the annotations.

